# A multi-component screen for feeding behaviour and nutritional status in *Drosophila* to interrogate mammalian appetite-related genes

**DOI:** 10.1101/2020.05.04.076489

**Authors:** J Chalmers, YCL Tung, CH Liu, CJ O’Kane, S O’Rahilly, GSH Yeo

## Abstract

More than 300 genetic variants have been robustly associated with measures of human adiposity. Highly penetrant mutations causing human obesity do so largely by disrupting satiety pathways in the brain and increasing food intake. Most of the common obesity-predisposing variants are in, or near, genes that are expressed highly in the brain, but little is known about their function. Exploring the biology of these genes at scale in mammalian systems is challenging. We therefore sought to establish and validate the use of a multicomponent screen for feeding behaviour and nutrient status taking advantage of the tractable model organism *Drosophila melanogaster*. We validated our screen by demonstrating its ability to distinguish the effect of disrupting neuronal expression of four genes known to influence energy balance in flies from ten control genes. We then used our screen to interrogate two genetic data sets. Firstly, we investigated 53 genes that have been implicated in energy homeostasis by human genome wide association studies (GWASs): of the 53 *Drosophila* orthologues studied, we found that 16 significantly influenced feeding behaviour or nutrient status. Secondly, we looked at genes which are expressed and nutritionally responsive in specific populations of hypothalamic neurons involved in feeding/fasting (POMC and AgRP neurons): 50 *Drosophila* orthologues of 47 murine genes were studied, and 10 found by our screen to influence feeding behaviour or nutrient status in flies. In conclusion, *Drosophila* provide a valuable model system for high throughput interrogation of genes implicated in feeding behaviour and obesity in mammals.

**Author Summary:** New high-throughput technologies have resulted in large numbers of candidate genes that are potentially involved in the control of food intake and body-weight, many of which are highly expressed in the brain. How, though, are we to find the functionally most relevant genes from these increasingly long lists? Appetite needs to be explored in context of a whole animal, but studies in humans and mice take a long time and are expensive. The fruit fly, while clearly evolutionarily distant, shares a surprising amount of biology with mammals, with 75% of genes involved in inherited human diseases having an equivalent in flies. In particular, the fruit fly has surprisingly conserved neuronal circuitry when it comes to food intake. Here we have developed a suite of four different functional assays for studying the feeding behaviour and energy balance in flies. We then used these assays to explore the effects of disrupting the expression of genes in the neurons of flies, that either are implicated in body weight through human genetic studies or are expressed and nutritionally responsive in specific populations of neurons involved in feeding. We show that the use of fruit flies are time and cost efficient, and are a valuable model system for studying genes implicated in feeding behaviour and obesity in mammals.

## Introduction

Obesity is arguably the greatest public health threat of the 21^st^ century (1), and is associated with co-morbidities such as type 2 diabetes, cardiovascular disease, hypertension and certain cancers (2). Whilst our changing lifestyle has undoubtedly driven the increase in obesity, there is a large variation in peoples’ response to this ‘obesogenic’ environment (3). Underlying this variable response is a powerful genetic element, with twin and adoption studies revealing the heritability of fat mass to be between 40% and 70% (4, 5).

Over the past two decades, a number of genetic and ‘omics’ approaches have been used to characterise the molecular and physiological mechanisms of food intake and body weight control. For instance, studies of human and mouse genetics have uncovered a number of circuits within the brain which play a central role in modulating mammalian appetitive behaviour (6, 7). The best characterised is the hypothalamic leptin-melanocortin signalling pathway, genetic disruption of which causes the majority of monogenic severe obesity disorders in both mice and humans (8–11). In addition, genome-wide association studies (GWASs) have, to date, identified more than 300 human genetic loci associated with variations in body mass index (BMI) (12, 13). The genes closest to these loci, which include many components of the melanocortin pathway, are primarily expressed in the central nervous system (CNS) (13, 14), and where their function is known, appear to influence food intake (6–11, 15–19).

In addition to genetic approaches, transcriptomic analyses of discrete neuronal populations are playing an increasingly important role in illuminating novel genes and pathways that may play a role in appetite control (20, 21). We know of at least two populations of neurons that sense peripheral nutritional signals and play a central role in the melanocortin pathway: POMC neurons decrease food intake when activated, while AgRP neurons increase food intake (22, 23), and there are a number of transcripts that are reciprocally regulated in these two populations (21). However, the function of most genes linked to obesity and appetitive behaviour remains unclear. One reason for the disappointing rate of translating the genetic signals into insightful biological knowledge is that investigations of candidate genes have to date been addressed in complex model organisms such as mice (24–32). Given the large resources required to generate and phenotype each murine model, they are not ideal for studying the effects of disruption of gene function at scale. There is, therefore, a requirement for a high throughput model to ‘pre-screen’ genes for a potential role in feeding behaviour, before committing the resources needed for more intensive studies of these candidates in mammalian models.

The fruit fly *Drosophila melanogaster* has generation time of only 10 days, a rate of up to 100 eggs produced per female fly per day (33), and a 10,000 fold lower cost to maintain fly stocks compared to mice (34). In addition, mutant fly lines and other resources are abundant and freely available within the scientific community (34). Although there clearly will be mammalian biology that is not replicated in flies, ~75% of genes involved in inherited human diseases have an orthologue in flies (35, 36). Furthermore, there is substantial conservation of tissue-specific patterns of gene expression, suggesting that there is also likely to be high levels of functional conservation (36).

Here, we report a high throughput *in vivo* functional screen for genes involved in mammalian feeding behaviour and nutritional status utilising *Drosophila*. Because feeding behaviour is largely controlled by the brain, we chose a neuron-specific approach involving RNA interference (RNAi) knockdown. We validated a suite of four assays by their ability to detect nutritional perturbation in wild-type flies, and to distinguish between positive and negative control genes. We then used these assays to screen two different genetic data sets. First, we studied 53 orthologues of human genes which are near single nucleotide polymorphisms (SNPs) that have been identified by GWAS to be robustly associated with human adiposity. In addition, we studied 50 *Drosophila* genes corresponding to 47 murine genes that are reciprocally regulated in POMC and AgRP neurons in response to an overnight fast. With both data sets, we demonstrate the utility of the tractable model organism in a high throughput genetic screen for food intake and body weight phenotypes, and identify candidates to pursue in further studies.

## Results

### Selection of assays for the high throughput screen

In order to create a *Drosophila*-based functional screen for appetitive behaviour and related traits, we selected four different assays: two that assessed energy intake (the CAFE assay and the fasting-induced feeding assay) and two that measured nutrient status (body mass and glucose levels). To validate these assays, we tested each for their ability to differentiate wild-type flies on a normal diet from those either fasted for 24 hours or placed on a high fat diet (HFD) for five days (Figure 1).

**Figure 1.**
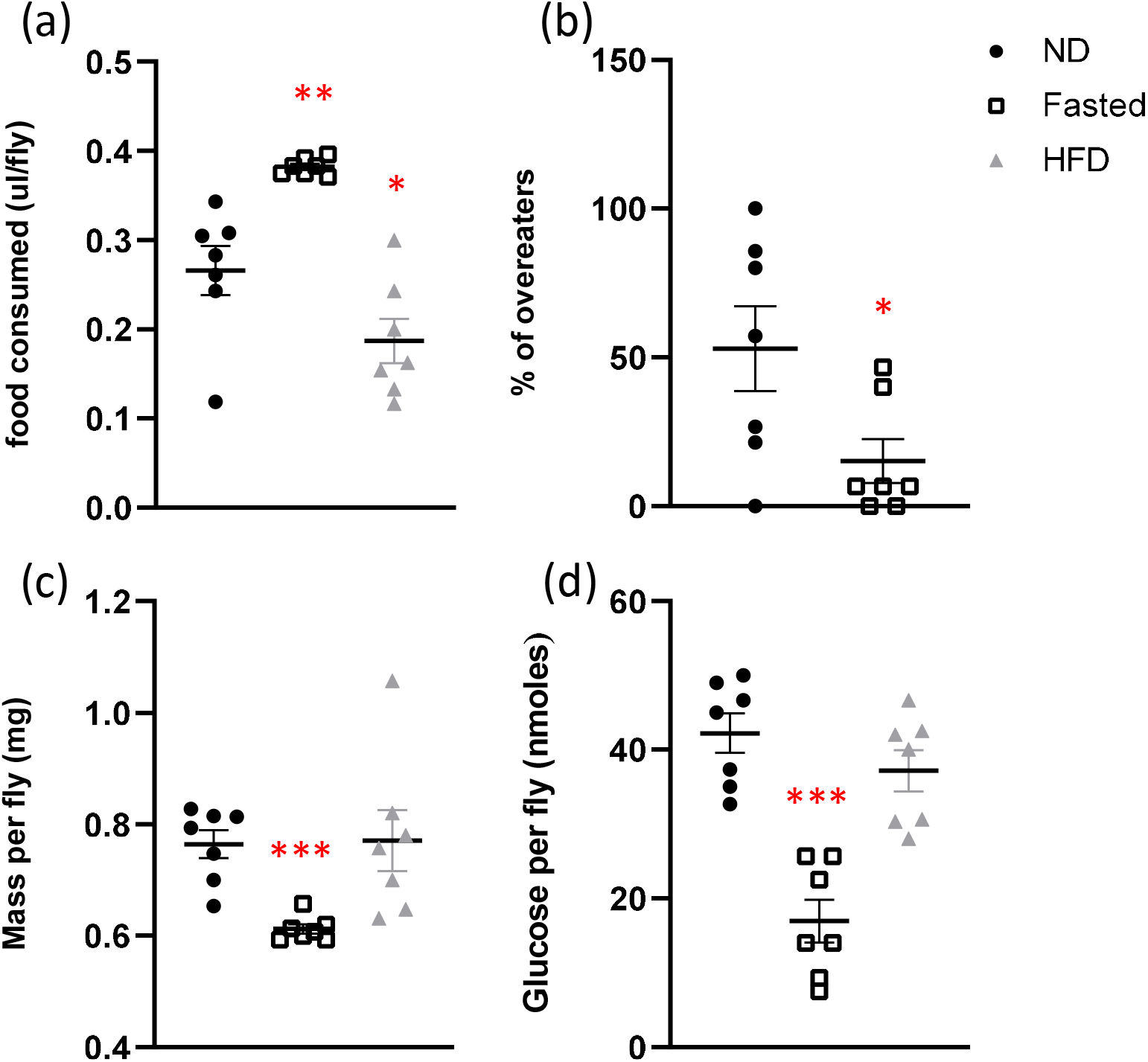
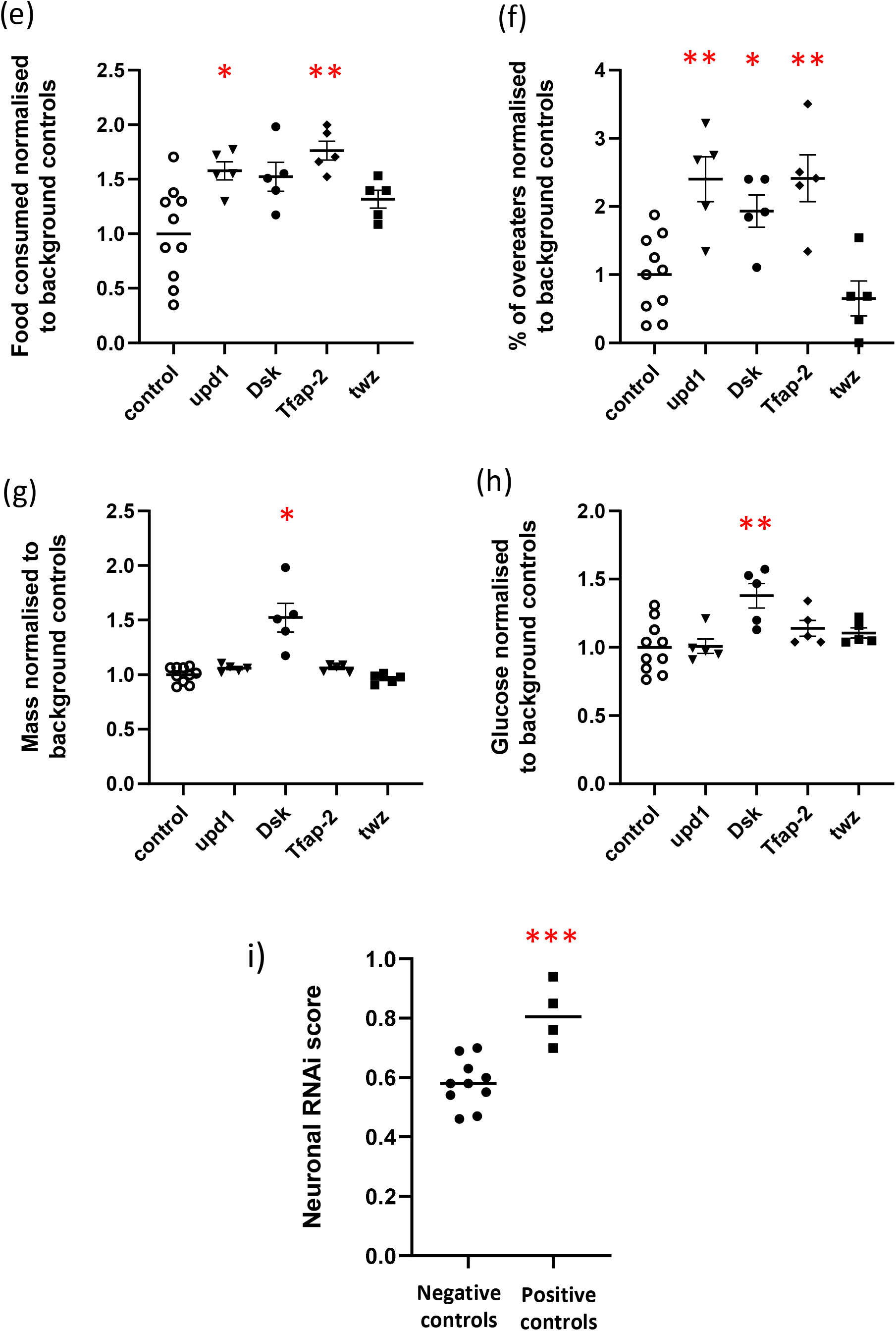
Validation of assays used in the functional screening. All four assays were able to distinguish flies fully fed on normal diet (ND) from flies fasted for 24hours (Fasted) or 5 days HFD (HFD). The behavioural assays were (a) the CAFÉ assay, to assess total food intake in a group of 8 flies over 24hours; and (b) the overfeeding dye ingestion assay, where ingested dyed-food was visually detected in the abdomen of adult flies after 20min fed *ad-lib* on non-dyed food following a 24hr fast, before being switched over to dyed food for 15min. The percentage of flies with coloured abdomens was calculated. The physiology assays were average measurement of 15 flies for (c) wet mass and (d) glucose storage. These assays were also able to distinguish flies with neuronal *elav-GAL4* driven knockdown on genes known to be involved in obesity, *albeit* not all four positive control genes showed a difference in all assays (e-h) as expressed relative to its respective controls. The overall phenotype score of all control genes presented in Table 1 showed a significant difference between the positive and the negative controls. Individual symbols indicates measurements from at least 5 repeats and lines denote the Mean ± SEM values among biological repeats. Statistical significance for the effect was determined using Student’s t-test (* p≤0.05, ** p≤0.01 or *** p≤0.001).

The CAFE assay measures the amount of liquid food (sucrose plus yeast extract) that flies consume from a capillary tube by tracking the distance moved by the meniscus over a given time. As expected, fasted flies consumed significantly more than the controls (144%; p=0.0012), while flies on a HFD consumed less (70%; p=0.0538; Figure 1a). The second assay measures fasting-induced food intake. Flies are fasted for 24 hours, then exposed to normal food for 20 minutes, before being transferred onto food coloured with dye for 15 minutes, which is then visible through their translucent abdomen. In the HFD group the number of flies with green stomachs decreased to 29% of controls (Fig 1b). In assays of nutrient status, fasted flies, as expected, had decreased wet mass (80%, p<0.0001) (Fig 1c) and glucose levels (40%, p<0.0001) (Fig 1d) when compared to flies fed *ad libitum*, while a HFD had no measurable effect on these parameters. Thus, the four assays were able to detect physiologically meaningful responses of wild-type flies to nutritional perturbation.

**Table 1.**
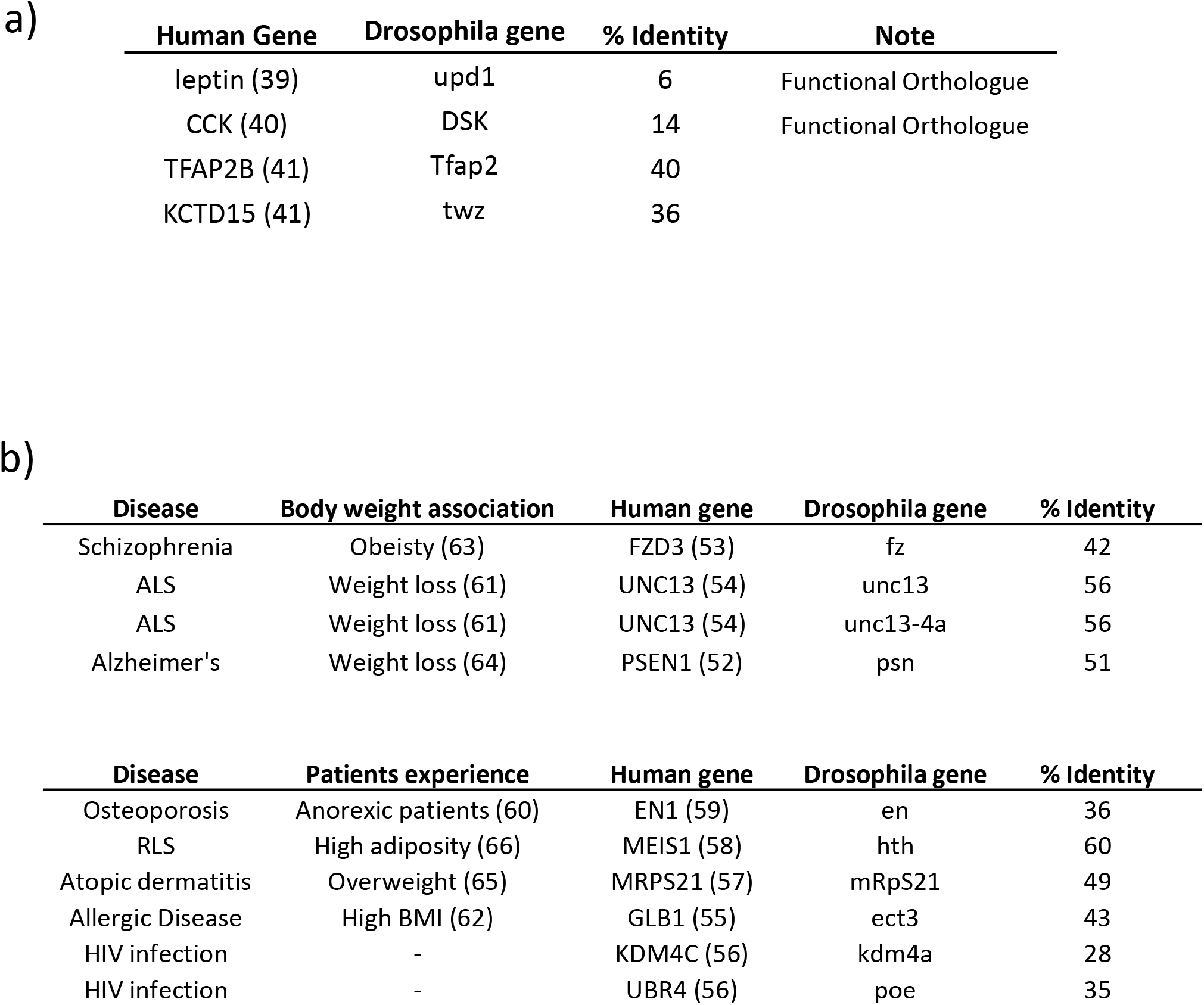
Positive and negative control genes. Positive and negative control genes were selected from the published literature. (a) The positive control genes have been shown to be involved in the neuronal control of feeding in mammals and all have been shown to have a phenotype in at least one of the screen assays. For the negative controls, some have a neuronal aetiology (b) and the others not with a strong neuronal association (c) but with primary symptoms that are non-metabolic.

### Validation of the high throughput screen using “control” genes

Next, we tested our screen on four genes known to play a role in *Drosophila* energy homeostasis (Table 1a). As most of the genes which have been linked to BMI are enriched for expression or function within the CNS (13), we took advantage of *the UASG-GAL4* system (37) to knockdown expression of each gene using RNAi specifically in *Drosophila* neurons using the neuronal driver *elav*-*GAL4* (38). Results from the neuronal RNAi were compared to background-matched control lines (either *KK-elavGAL4*, *GD-elavGAL4* or *KKtiptoe-elavGAL4* as appropriate; Supplementary Table 6). As one example, neuronal knockdown of *upd1*, a *Drosophila* orthologue of leptin (39), led to flies ingesting 1.5x more in the CAFE assay (p=0.0149; Figure 1e) and 2.4x more flies ingesting dyed food in the fasting-induced feeding assay (p=0.0013; Figure 1f).

To take advantage of the multiple assays, we used a simple algorithm to produce an integrated score that reflects the overall phenotype resulting from neuronal knockdown of each gene. The score is calculated by taking the sum of the p-values from the four assays for each gene, and subtracting it from 1. Thus, the more significant the results for a given gene, the lower the p-value, and the closer the score would be to 1. Using this method produced an integrated score of 0.76 for *upd1*. The scores for the other positive controls were 0.85 for *DSK* (the *Drosophila* orthologue of *CCK;* (40), 0.94 for *Tfap2* and 0.70 for *twz* (the orthologue of *TfAP-2* and *KCTD15,* respectively (41).

In addition, we tested 10 ‘negative’ control genes that have been linked to diseases other than obesity (Table 1b). When the scores for all of the controls were plotted, the flies with neuronal knockdown of ‘positive’ control genes had a significantly higher average score than the ‘negative’ controls (p=0.0006; Figure 1i). Based on its ability to distinguish these two groups, we set a score of 0.80 (an average of p=0.05 for each assay) as a threshold for selection of genes of interest, with consideration given to genes with a score ≥0.75.

### Screening of candidate genes from GWASs of BMI

Having validated our suite of assays, we used them to screen a number of candidate genes highlighted by GWAS to be associated with BMI. We focussed on the 36 genetic loci identified by Speliotes et al. (42) and Lu et al.(43). Our workflow for identifying the related *Drosophila* genes is summarised in Figure 2. We chose to study the closest gene to each of the 36 BMI-associated SNPs (Supplementary Table 1), as well as additional genes within 500 kb in the vicinity of seven of the SNPs, giving a total of 53 human genes (Supplementary Tables 1 & 2). 43 of the human genes (81%) had at least one *Drosophila* orthologue, and some had more than one, giving a total of 56 *Drosophila* genes (Figure 2). RNAi lines were available to study for 53 of these genes.

**Figure 2.**
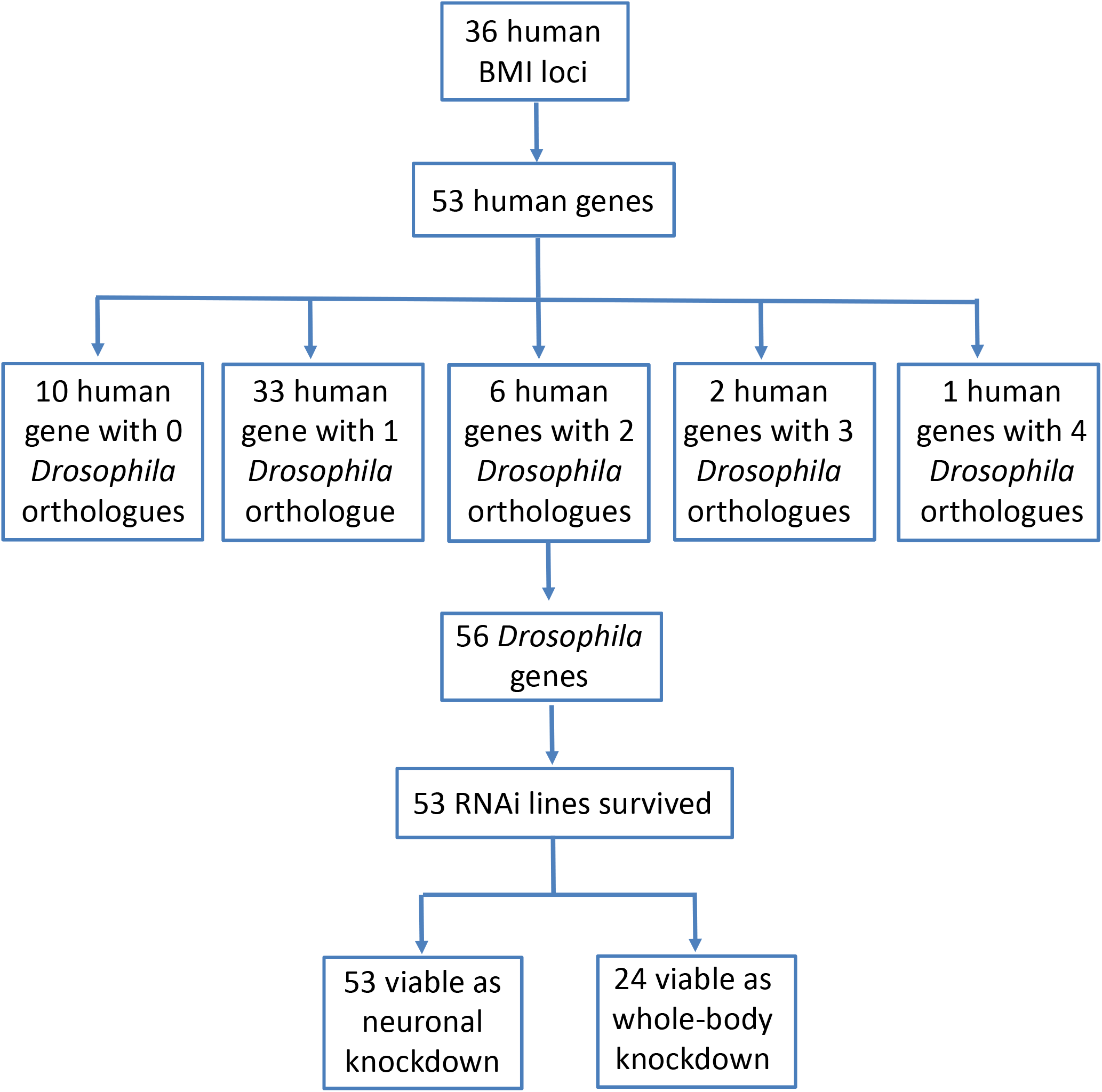
Selection of the BMI GWAS gene. Summary of the experimental design of the *Drosophila* functional screen of candidate obesity gene based on GWAS BMI association loci. All genes had at least one orthologue in *Drosophila*, resulting in the study of 53 potential candidate gene for obesity.

When we generated whole-body RNAi knockdowns (using the *Act5C-GAL4* driver), 29 of the 53 genes (55%) were not viable as adults (Figure 3). This is much higher than the genome-wide figure of 25% (36), suggesting that the BMI-associated GWAS genes are genes enriched in genes which are essential for life.

**Figure 3.**
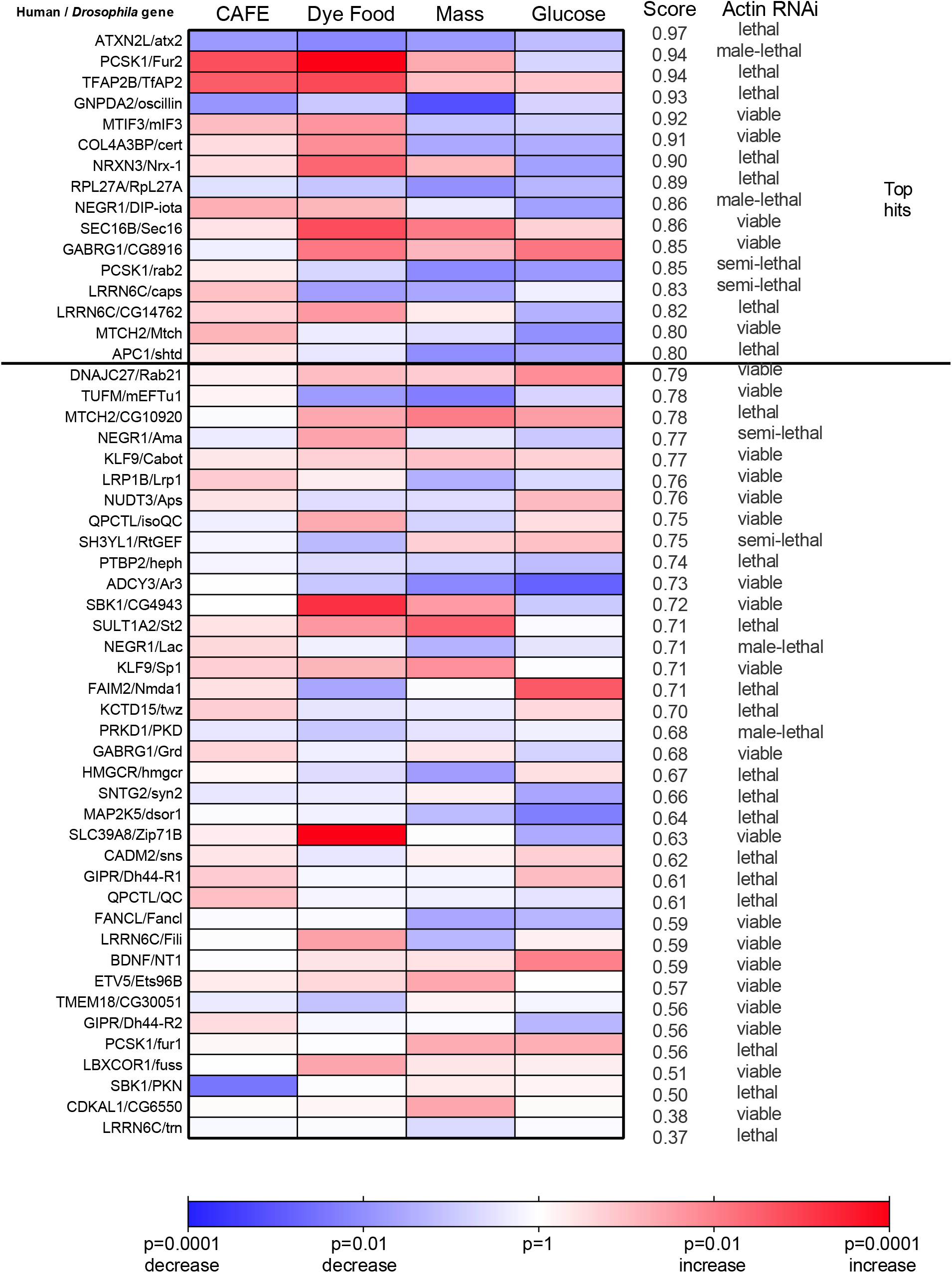
Neuronal knockdown of BMI GWAS genes. Summary of the results from the functional scan of the 53 BMI GWAS loci with *Drosophila* orthologues, visualized as a heatmap. Results compare neuronal-specific knockdown, using the drivers *elav-GAL4*, compared to the respective control RNAi-lines. Colour corresponds to significance determined by Student’s t-test for each assay; and the ‘score’ is calculated by summing up the four p-values from each assay and subtracting it from 1, thus the closer the number to 1, the more significant the results. Genes are ranked by score and we set ≥ 0.80 (an average of p=0.05 for each assay) as the threshold for a ‘hit’. The viability of flies with whole body knockdown of the gene using *actin-GAL4* are shown on the far right-hand column.

As above, we used *elav-GAL4* to perform neuron-specific knockdown of each of our study genes. In contrast to the whole-body knockdowns, neuronal RNAi of all 53 genes produced viable adults. The phenotype results for each gene are summarised in Figure 3 ranked by score. Neuronal knockdown of 16 of the fly genes, corresponding to 14 human genes, resulted in a phenotype score of 0.80 or more in our assays. It is interesting to note that 53 of the 56 *Drosophila* genes selected for our screen are expressed in the *Drosophila* brain (44, 45), suggesting conservation of expression. The three genes which are not neuronally expressed (*Grd/GABRG1*, *CG10920/MTCH2* and *Ets96B/ETV5*), should therefore be unaffected by neuronal RNAi, all scored less than 0.80.

Several genes with a score above 0.8 in our *Drosophila* assays have mouse orthologues that play a role in energy homeostasis (*ATXN2L, PCSK1, NEGR1, MTIF3, SEC16B*), increasing our confidence that our screen can reveal biologically plausible candidates. Other genes (*GNPDA2, NRXN3, GABRG1, LRRN6C, APC1*) are relatively unexplored in relation to energy homeostasis, are relatively unexplored in relation to energy homeostasis, marking them as prime candidates for further study.

At seven of the GWAS loci, multiple human genes were studied (Supplementary Table 2). At three of these loci, the nearest gene to the SNP resulted in the highest score compared to other genes in the vicinity: *GNPDA2 (0.93), RBJ* (0.79) and *QPCTL (0.75)*. At four other loci, the highest score was observed by neuronal knockdown of ‘non-closest’ genes: *ATXN2L* (0.97), *COL4A3BP* (0.91) and *APC1* (0.80). In addition, at the rs16951275 locus, neither the nearest gene (*LBXCOR1*) nor the nearby gene (*MAP2K5*) had a score >0.70, perhaps implicating other genes further than 500 kb away.

Baranski and colleagues (46) previously performed a screen of triglyceride levels in *Drosophila* where BMI GWAS candidate genes were knocked down using RNAi in both the CNS and fat body concurrently. There were 16 *Drosophila* genes in common between their screen and ours (Supplementary Table 3). In their screen, five of these genes showed an increase in triglycerides when knocked down. In our screen, three of these (*NRXN3*, *SEC16B*, and *COL4A3BP*) had scores >0.80, and the other two (*NUDT3* and *SBK1*) had scores >0.70 (Figure 3; Supplementary Table 3). Interestingly, *Atx2* (corresponding to the human gene *ATXN2L*) showed no phenotype in the triglyceride screen, but achieved the highest score of 0.97 in our screen. However, in our hands, the knockdown of *Atx2* resulted in flies that ate less, whereas mice lacking *Atxn2* show a phenotype of adult-onset obesity (47). So while perturbation of its expression resulted in opposite feeding phenotypes in flies and mice, it is clear *Atx2* and *Atxn2* play a role in energy balance. Interestingly, when Baranski and colleagues knocked down RPL27A within the CNS and fat body, it resulted in a lethal phenotype. In our hands, neuronal specific manipulation of RPL27A expression resulted in a score of 0.89, thus implicating RPL27A in the control of food intake.

### Screening of genes which are reciprocally regulated in mammalian POMC and AgRP neurons

Next, we turned our attention to candidate genes emerging from transcriptomic analyses of key neurons known to play a role in the control of mammalian appetitive behaviour. Henry et al. (21) reported the expression levels of 35,266 mouse genes in both POMC and AgRP neurons. We utilised this publicly available dataset to select a list of candidate transcripts with potential relevance to feeding. Our workflow is summarised in Figure 4. Briefly, of the 1,038 and 3,554 genes whose expression significantly changes (>1.5x; p<0.05) upon fasting in POMC and AgRP neurons, respectively, 192 respond in a reciprocal manner. Using ENSEMBL, we identified *Drosophila* orthologues for 58% of these mouse genes, including some with multiple orthologues, and two mouse genes with the same fly orthologue. Overall, this resulted in a list of 157 *Drosophila* genes. To maximise the likelihood of generating relevant biological insights, we selected only genes with sequence homology >30% and which are expressed in the *Drosophila* CNS either at the adult or larval stage (44). This resulted in inclusion of 61 genes in our screen (Figure 4). As with the GWAS genes, lines enabling RNAi knockdown of each of these *Drosophila* genes were identified. For two genes no RNAi lines were available, and for another six the only lines available had multiple off-target effects so were excluded from further analysis. In total, this screen assayed 11 ‘orexigenic’ genes (decreased expression in POMC neurons and increased expression in AgRP neurons during a fast) and 39 ‘anorexigenic’ genes (increased expression in POMC and decreased expression in AgRP neurons during fasting) (Figure 5; Supplementary Table 4). Once again, we utilised *elav-GAL4* to knockdown these genes specifically in *Drosophila* neurons.

**Figure 4.**
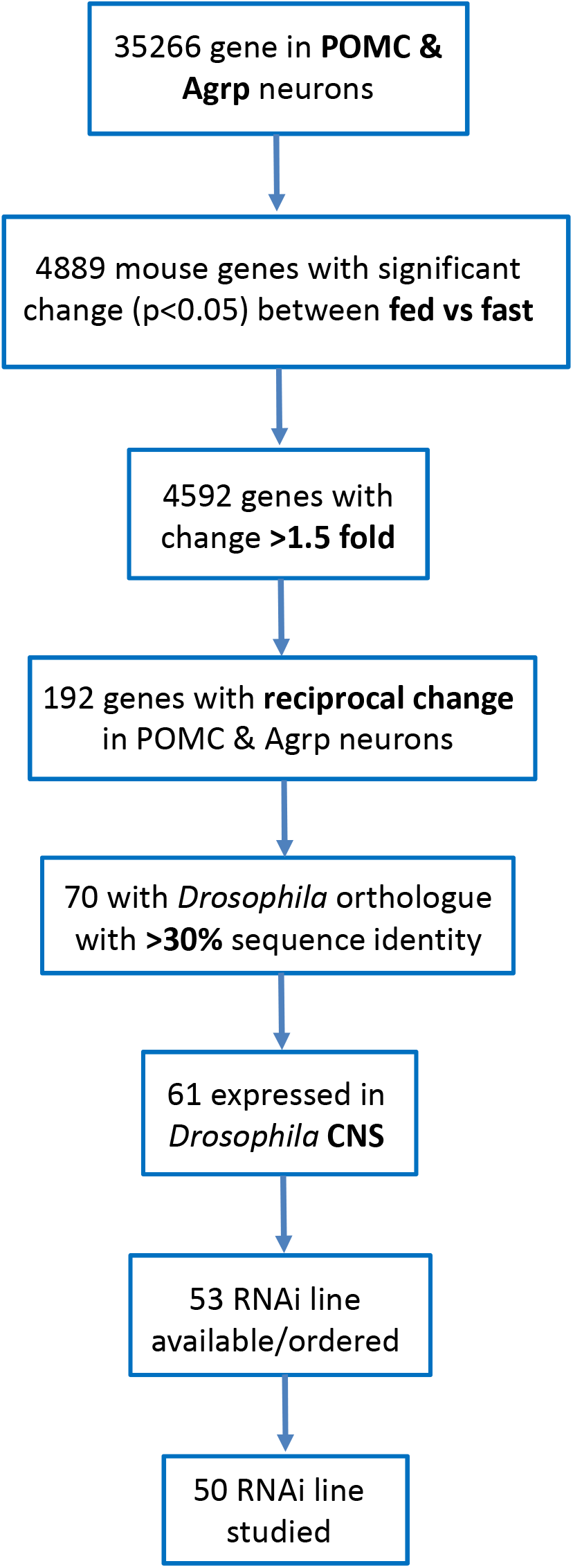
Selection of genes reciprocally regulated in murine POMC and AgRP neurons in response to fast. Workflow of the obesity and anti-obesity candidate gene selection scheme. Candidate genes were identified from Henry et al. after *in silico* pre-screening based on the significance of the changes in a reciprocal fashion in the POMC and AgRP neurons in response to a fast. The threshold was set to be at least a 1.5 fold change in expression with a p<0.05.

**Figure 5.**
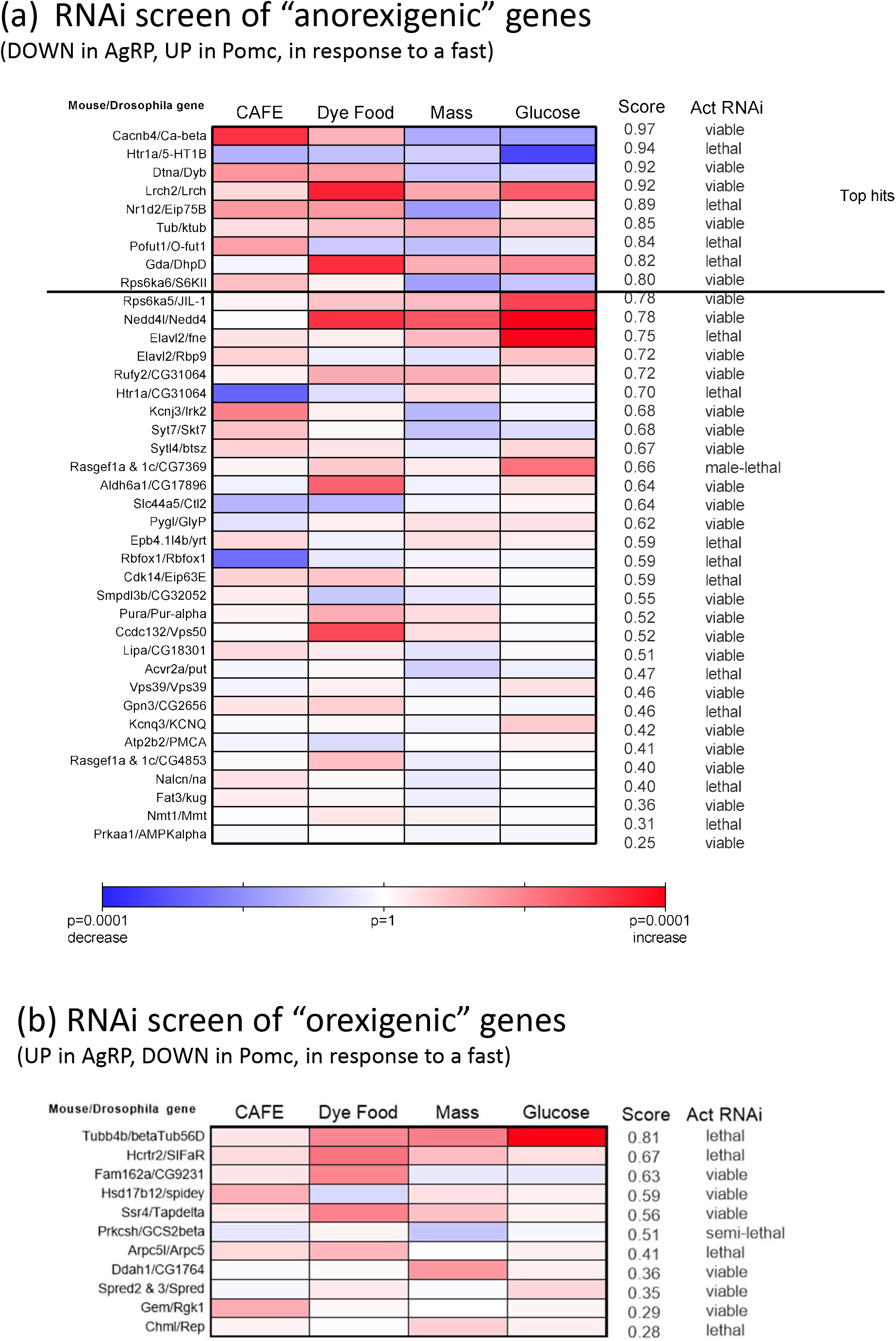
RNAi screen of murine “anorexigenic” and “orexigenic” genes. Summary of phenotypes and ranked overall score of flies with neuronal RNAi knockdown of the murine anorexigenic genes (a) and orexigenic genes (b). Genes were ordered accordingly to the overall score and the top hit genes are those with score ≥0.8. The viability of flies with whole body knockdown of the gene using *actin-GAL4* are shown on the far right column.

The results of the screen, with genes ranked by score, are shown in Figure 5. The scores have a larger range (0.25 to 0.97) than was seen with the GWAS genes. Nine of the ‘anorexigenic’ genes (*Cacnb4, Htr1a, Dtna, Lrch2, Nr1d2, Tub, Pofut1, Gda, Rps6ka6*) and one ‘orexigenic’ gene (*Tubb4b*) had a score ≥0.80.

For all genes, we additionally collected triglyceride level data, and showed that the wet mass is a reliable substitute for triglyceride measurement as the overall selected candidate gene list remained the same with either measurement (Supplementary Table 5). However, it is worth noting that it is easier and quicker to measure mass than triglyceride levels.

For two of the genes with a score >0.80 (*Htr1a*, *Tub*) and six of the genes with a score between 0.70 and 0.80 (*RPS6ka5, Nedd4l, Elavl2, Rufy2* and *Htr1a*), there is published evidence from mouse and/or human studies for a role in energy homeostasis.

## Discussion

The aim of this study was to develop an *in vivo* high-throughput screen of feeding behaviour and energy balance phenotypes using *Drosophila* as a model. We then successfully demonstrated its utility by using it to study genes identified by GWAS to be associated with BMI (14, 42), as well as genes reciprocally regulated by fasting in Agrp and POMC neurons that play a key role in the central melanocortin pathway (21).

Previous *Drosophila* screens have focussed on single metabolic measures in either adult flies (46, 48–50) or larvae (51). In contrast, we have pulled together a suite of 4 assays that measured different aspects of energy homeostasis, including food intake, body weight and glucose levels. Using multiple assays to generate an integrated score has two major advantages; firstly, it is less likely to give false positive results; and secondly, it increases the sensitivity of the screen, allowing genes that do not show a large phenotype in any single assay, but which have subtle changes in multiple assays to be highlighted. An issue we initially grappled with was our selection of ‘negative controls’, which fell into two categories; those genes that are associated with neuronal diseases (52–54) and those associated with diseases of the periphery (55–59). The problem is that many human diseases are associated with changes in body weight (60–66), as either a cause or a consequence (67). The challenge was therefore differentiating these genes from those that primarily influenced feeding behaviour. While the results were not black or white, our suite of assays were able to discriminate the negative from the positive controls (Figure 1), with average scores significantly different between the two groups. It did mean that the scoring threshold, above which we would consider taking genes forward for further study, was set quite high, at 0.8. However, since the principle aim of this screen was to prioritise candidate genes to follow up, our goal was to avoid false positives where possible, and we were comfortable with this high threshold.

Several of the human genes that were highlighted by our screen (*PCSK1, ATXN2, NEGR1, MTIF3* and *SEC16B*) have already been shown to play a role in energy homeostasis in mouse models (24, 27, 28, 30, 32), demonstrating that the screen is effective and produces biologically relevant results. Critically, we also identified genes that are relatively unexplored in relation to energy homeostasis, marking them as prime candidates for further study. These include *GNPDA2, NRXN3, GABRG1, LRRN6C, APC1, COL4A3BP, RPL27A and DNAJC27*. *NRXN3* (Neurexin 3) for example, is a neuronal cell surface protein involved in cell recognition and cell adhesion (68); *GABRG1* (GABA(A) Receptor Subunit Gamma-1) belongs to the ligand-gated ionic channel family and predominantly expressed in the brain reward circuitry and may be implicated in addiction (69); while *GNPDA2* is an enzyme that catalyses the deamination of the glucosamine-6-phosphate shown to be a critical gene for lipid and glucose metabolism in human adipose-derived mesenchymal stem cells (70). The role that these genes play in the regulation of feeding behaviour remains to be uncovered. Some genes, including *Nr1d2* and *Cacnb4,* showed a *Drosophila* phenotype of significantly increased feeding, but decreased mass and/or glucose levels, suggest increased metabolic demands. We are unable to explain these apparently discordant phenotypes without detailed follow-up studies, but *Nr1d2* has been shown to circadian rhythm and could potentially play a role in energy homeostasis (71).

The advantages of using multiple assays were demonstrated when we compared our results to the recent study by Baranski and colleagues, who also studied GWAS identified genes associated with BMI by knocking them down in *Drosophila* (46). There were 16 genes in common between both our studies, 5 of which were identified by Baranski and colleagues to have increased triglycerides storage when knocked down in the brain and fat body. 3 of these 5 genes, *NRXN3*, *SEC16B* and *COL4A3BP* were positively identified by our screen with a stringent cut off of overall score >0.8, with all 5 resulting in a score >0.7, providing independent validation of the utility of our approach. Crucially, our screen identified *ATXN2L,* which although when knocked down did not result in an increased triglyceride storage, came up positive using our screen due to its consistent anorexigenic effect on food intake. Evidently, mice lacking *Atxn2* are sensitive to diet induced obesity (47) and evidence suggested that Atxn2 modulates nutrition and metabolism by regulating the pathways for the metabolism of branched-chain and other amino acids, metabolism of fatty acids, and the citric acid cycle (72). While we agree that triglyceride levels are an important readout of nutrient status, we find the use of body mass to be a useful proxy for use within a high-throughput screen (Supplementary Table 5).

Another unique element to our study was our decision to use a neuron-specific RNAi approach. Genes identified by GWAS to be associated with BMI are enriched in the CNS (13), and 53 of the 56 genes that we studied were expressed in the *Drosophila* CNS. When knocked down in the whole body, 55% of the lines were lethal. This is a far higher proportion of lethality as compared to when we knocked down fasting related genes (38%) and to the *Drosophila* genome-wide figure of 25% (36), suggesting these genes are critical to viability. Even in the Baranski study, where they knocked down the genes in the brain and fat body, they reported 13% lethality. In contrast, all of our neuronal specific knock-down lines were viable. So our more tissue-targeted approach allowed us to screen more genes and obtain more phenotypic information.

A perennial problem with GWAS is that the vast majority of SNPs associated with disease are located in non-coding regions, making the identification of the ‘causative’ gene(s) driving the phenotype challenging. We used our high-throughput screen to assay multiple genes at 7 of the BMI loci, to see this could shed light on the potential causative gene. Of the 24 human genes studied in these 7 loci, 6 were scored as a ‘positive hit’ by our screen: *ACP1* (near rs2867125), *RBJ* (near rs713586), *GNPDA2* and *GABRG1* (near rs10938397), COL4A3BP (near rs2112347), and *ATXN2L* (near rs7359397). *ATXN2L, ACP1, COL4A3BP* in particular are not the genes closest to their respective SNPs. This is comparable to the results reported by Baranski, where in 16% of cases, knocking down the ‘non-nearest gene’ gave the largest phenotype.

Of course, the major limitation of working with *Drosophila* is that it is only distantly related to humans. However, given that energy homeostasis and feeding are important fundamental processes for all organisms, it is likely that there will be conservation of circuitry and pathways between species. In fact, nearly 80% of the human BMI genes and a similar percentage of the murine hypothalamic fasting genes had a *Drosophila* orthologue.

Once the four assays were established, we were able to screen around 60 different *Drosophila* lines in 6 weeks, at a cost of less than £60 per gene, resulting in 30% of human genes screened being highlighted as warranting further study. In conclusion, we demonstrate that the use of *Drosophila* for screening feeding behaviour and energy balance phenotypes is effective in moving from large lists obtained from whole genome or transcriptomic approaches, to more discrete lists of relevant genes, where ‘lower throughout’ and more time-consuming functional analyses in mammalian models can be focussed on. Furthermore, the detrimental effects seen in many of the whole-body vs neuronal specific *Drosophila* knock-downs suggest that, at least for a subset of the candidate genes, neuronal specific models may also be needed to explore genotype/phenotype relationships relevant to energy balance in higher organisms

## Materials and Methods

### Fly husbandry

All flies were raised on Normal diet (ND: agar, dextrose, maize, yeast, nipagin, water). Some assays required use of dye food (ND + 1% Fast Green FCF dye (Sigma) or high fat diet (HFD: ND + 20% coconut oil). Experimental flies were maintained at 25°C and 60-70% humidity in a 14 hour light: 10 hour dark cycle, except for flies on HFD which were kept at 20°C in the dark. Unless otherwise noted, we used male flies that were between 5-10 days old for the experiments and 15 flies per sample, 5 biological replicates per genotype.

*Drosophila* orthologues of the human genes of interest were identified using the ENSEMBL (73) or NCBI BLAST searches of publicly available protein sequences (74, 75). Fly stocks (listed in Supplementary Table 6) were acquired from the Vienna Drosophila Resource Center (VDRC) and the GAL4 lines from the Bloomington Drosophila Stock Center. For RNAi stocks, preference was given to GD lines to avoid potential complication with tiptoe gene expression (76). Two GAL4 lines were used here, *elav-GAL4* which expressed in neurons (38) and *act-GAL4* which is constitutively expressed throughout the body. UAS-RNAi lines were crossed to Gal4 lines (*elav-Gal4* or *act-Gal4*) and controls were created by crossing each Gal4 line to GD, KK and KK_tiptop_. Experimental flies were compared to the relevant control background (Supplementary Table 6). Three of the GD lines (V14, VV22 and VV46) were compound X stock from VDRC thus we crossed onto an FM7 balancer prior to phenotyping.

To standardise the effects of parental environment on offspring fitness, all UAS-siRNA stocks were kept in bottles at an approximately constant density. For the generation of flies for phenotyping, virgin female UAS-siRNA flies were crossed to male GAL4 flies and then the male offspring were collected. To standardise the effects of parental age on offspring fitness, crosses were set up using flies which were 1-5 days old. Briefly, on day 1, 5 *UAS-RNAi* female virgins were placed in a vial of ND at 25°C with 2 *elav-Gal4* or 5 *act-Gal4* males. On day 4, these parents were removed. On day 14, offspring were transferred to a new vial and allowed to mate, as mating alters gene expression and metabolic parameters (77). On day 15, the females were removed, and on day 16, male flies were placed on starvation media for 3 hours to synchronise their metabolism, before being returned to the appropriate diet for the experiment. As *Drosophila* activity and feeding behaviours are affected by age and circadian rhythms (78), all assays were done within a certain window of time relative to the synchronisation.

### PCR-based diagnostic assay to determine appropriate background line for the “KK” collections

Genomic DNA was isolated by crushing flies in squishing buffer (10mM Tris, pH8.2, 1mM EDTA, 25mM NaCl, 200ug/ml protease K), 30min incubation at 37^c^C, and inactivate the proteinase K by heating to 95°C for 5min. We used the PCR-based diagnostic assay to determine KK insertion site as described in Green et al. (76). Briefly, occupancy of the transgene insertion site at the annotated insertion reported by the VDRC, 40D, was determined in multiplex PCR. Following primers and the PCR using these primers yields a ~450bp product in the case of an insertion, or a ~1050bp product in the case of an empty insertion site, at 40D.

40D Genomic_F 5’- GCCCACTGTCAGCTCTCAAC -3’

pKC26_R 5’- TGTAAAACGACGGCCAGT -3’

pKC43_R 5’- TCGCTCGTTGCAGAATAGTCC-3’

For the insertion at the non-annotated insertion, 30B (thus use Tiptoe as control), the pKC and pKC43R primers were multiplexed with the following primer

30B_Genomic_F 5’- GCTGGCGAACTGTCAATCAC -3’

PCR using these primers results in a ~600bp product in the case of an insertion, and a _1200bp product in the case of an empty insertion site, at 30B.

PCR was perfomed with the following programme: 95 ^c^C, 120s initial denaturation, 30 cycle reaction (95 ^c^C 15s denaturation, 50 ^c^C 15s annealing, 72 ^c^C 45s extension), and a final 72 ^c^C, 120s extension. These primers amplify a 700bp product within empty pKC26, and do not amplify if a RNA hairpin-encoding sequence is inserted PCR was performed.

### CAFE assay

The Capillary Feeder (CAFE) assay is an accurate method of measuring food intake in *Drosophila* (79) where the liquid food in the calibrated capillariy glass tubes (5ul, VWR Internation) are the only food available to the flies under study. The CAFE assay was performed with a custom-made acrylic cap with four holes that will hold 200ul pipette tips, two of which act as air holes and two hold the glass capillaries (VWR 53432-706). CAFE chambers were made from standard fly vials (25mm x 95mm) with 3ml of 1% agar that serves as a water source and maintains internal chamber humidity. Eight flies were placed onto each vial, two capillary tubes filled with liquid food contain 5% of sucrose and 5% of yeast extract via capillary action. The top of the meniscus was marked, and the filled capillaries were inserted into the chamber through the lid, and left at 25°C. The movement of the meniscus was measured and evaporation (as measured by a vial containing no flies) subtracted to give the volume of food eaten and 24h feeding is recorded after a day of habituation in the CAFE.

### Over-feeding dye assay

This protocol was modified from Williams et al. 2014 (46) and is particularly useful for examine the satiety signals involving in food intake. Fifteen male flies were fasted for 24 hr by keeping in the vials with 1% agar. On the day of the experiment, flies were transferred to normal diet and allowed to feed. After 20 minutes, the flies were then transferred to vials containing 1% Fast Green dye for 15 minutes. The number of flies with visible dye in their abdomen (mid-gut and/or crop) was counted under dissecting microscope and scored as a percentage of the total number in the vial.

### Wet mass & Gloucose assay

Groups of 15 *Drosophila* were frozen on dry ice and weighed using a microbalance (Sartorius), then stored at −80°C for glucose analysis. For glucose analysis, frozen *Drosophila* were placed in the FastPrep tubes containing Lysis Beads and Matrix D (MP Biomedicals) and 350μl of cold PBST (PBS + 0.05% Tween-20), then homogenised using a FastPrep-24 homogeniser (MP Biomedicals) for 60s at 6m/s. Solutions were centrifuged (16100rcf, 4°C, 3 minutes) to pellet debris, and 300μl of supernatant pipetted into a fresh Eppendorf on ice. Homogenates were heat-inactivated (5 minutes, 70°C), then the supernatant was re-centrifuged (16100rcf, 4°C, 3 minutes) to pellet debris, and ~100μl transferred into a fresh tube and stored at −80°C for glucose analysis. Glucose analysis was performed by the Cambridge Core Biochemical Assay Laboratory (www.cuh.nhs.uk/core-biochemical-assay-laboratory) using enzymatic assays. Glucose amounts were normalised to number of *Drosophila*.

### Triglyceride assays

For triglyceride analysis, *Drosophila* homogenate were prepared as described above for glucose analysis except the samples were not centrifuged prior to the analysis. The Cambridge Core Biocheical Assay Laboratory did triglyceride analysis using enzymatic assays.

### Scoring algorithm

The scoring system ‘sums up’ data for all four phenotypes done in adult flies in order to give a quantitative measure of the overall phenotype. The scoring takes an average of the p values for each assay, and subtracts from 1 so that more significant results give higher scores.

### Statistical Analysis

Each assay was repeated at least 5 times on independent crosses. Mean and standard error mean from all replicates of each experiment were calculated and analysised using GrapPad Prism. Data from all the assays was analysed using the unpaired homoscedastic Student t-test. ANOVA with appropriate post hoc analysis for multiple comparisons were employed where appropriate.

## Supporting Information Captions

**Supplementary Table 1.**
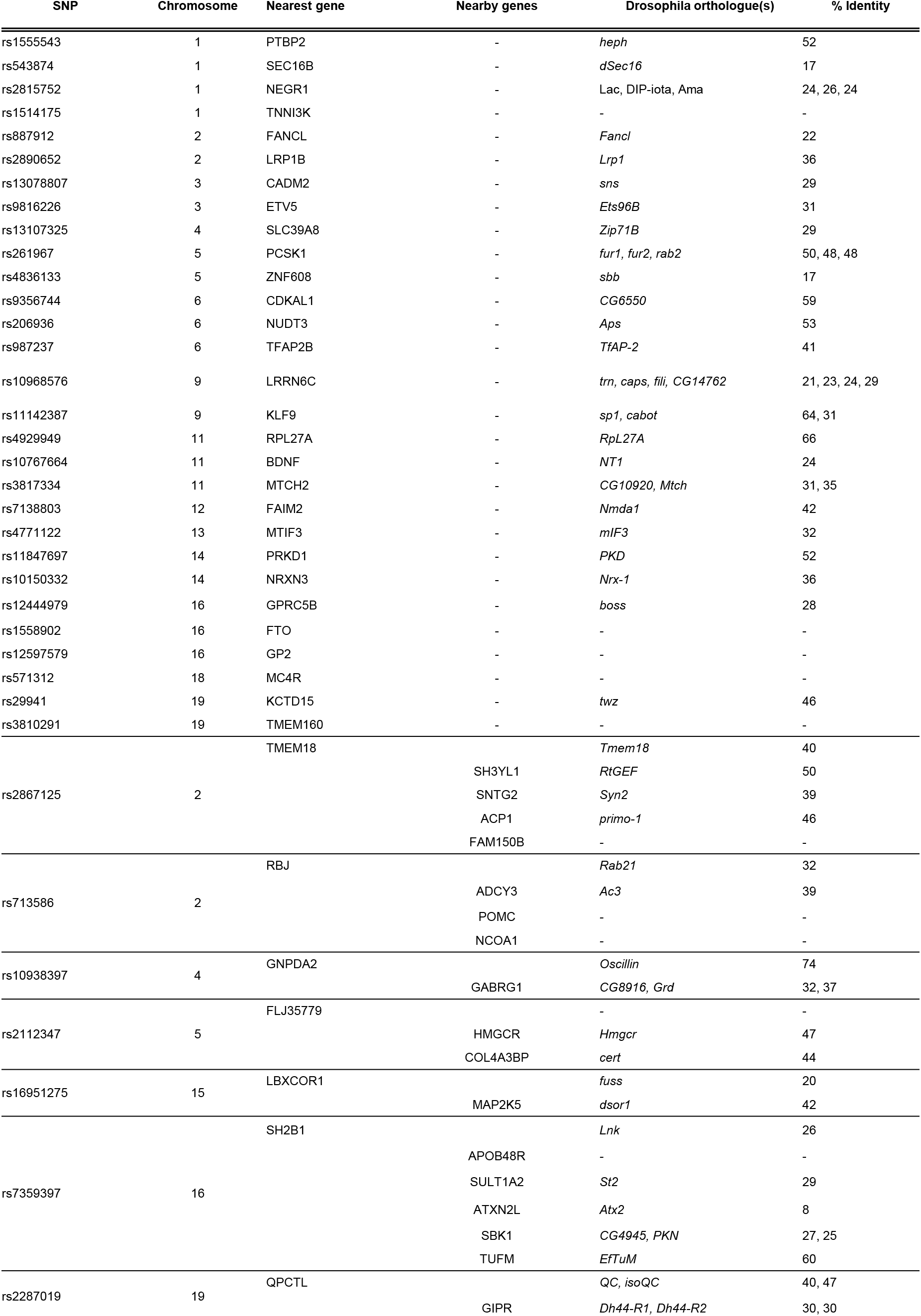
List of all human genes located near the 36 genome-wide significant loci on BMI and their respective *Drosophila* orthologues identified using the ENSEMBL or NCBI BLAST searches of protein sequences. SNP positions listed were based on Locke et al.

**Supplementary Table 2.** List of the multiple genes within the 500kb window vicinity of the seven SNPs identified in BMI GWAS loci that were included in the functional *Drosophila* screen. First row at each locus (in red) represents nearest gene, other rows represent the nearby genes in order of the calculated distance from the SNPs.

| SNP | Chromosome | Position | Notable genes | Gene Start | Gene end | Distance (kb) | Drosophila Orthologue | Score |
| --- | --- | --- | --- | --- | --- | --- | --- | --- |
| rs2867125 | 2 | 622,827 | *TMEM18(N)* | 663877 | 677406 | 41 | *CG30051* | 0.56 |
|  |  |  | FAM150B | 279558 | 288872 | -343 | - | - |
|  |  |  | ACP1 | 264869 | 278283 | -358 | *Shtd* | 0.8 |
|  |  |  | SH3YL1 | 218136 | 264866 | -405 | *RtGEF* | 0.75 |
|  |  |  | SNTG2 | 1087 | 50794 | -622 | *Syn2* | 0.66 |
| rs713586 | 2 | 24,935,139 | *RBJ/DNAJC27* | 24943636 | 24972094 | 8 | *Rab21* | 0.79 |
|  |  |  | ADCY3 | 24819169 | 24920237 | -116 | *Ar3* | 0.73 |
|  |  |  | POMC | 25160853 | 25168851 | 226 | - | - |
|  |  |  | NCOA1 | 24491254 | 24770702 | -444 | - | - |
| rs10938397 | 4 | 45,180,510 | *GNPDA2(N)* | 44701795 | 44726634 | -454 | *oscillin* | 0.93 |
|  |  |  | *GABRG1(B)* | 46035769 | 46124065 | 855 | *CG8916* | 0.85 |
| rs2112347 | 5 | 75,719,417 | *FLJ35779/POC5* | 75674124 | 75717488 | -45 | - | - |
|  |  |  | COL4A3BP | 75368486 | 75551981 | -351 | *cert* | 0.91 |
|  |  |  | HMGCR | 75336334 | 75362116 | -383 | *hmgcr* | 0.67 |
| rs16951275 | 15 | 67,784,830 | *LBXCOR1* | 67819704 | 67834561 | 35 | *fuss* | 0.64 |
|  |  |  | *MAP2K5(B,D,N)* | 67542606 | 67807117 | -242 | *dsor1* | 0.51 |
| rs7359397 | 16 | 28,874,338 | *SH2B1(B,M,Q)* | 28846600 | 28874213 | -28 | *Lnk* | (unwell) |
|  |  |  | *TUFM(Q)* | 28842411 | 28859562 | -32 | *mEFTu1* | 0.78 |
|  |  |  | *ATXN2L(Q)* | 28823048 | 28837237 | -51 | *atx2* | 0.97 |
|  |  |  | *SULT1A2(Q);* | 28591943 | 28597079 | -277 | *St2* | 0.71 |
|  |  |  | *APOBR(M,Q)* | 28494649 | 28498970 | -375 | - | - |
|  |  |  | *SBK1(Q,D)* | 28258687 | 28323849 | -550 | *CG4945* | 0.72 |
| rs2287019 | 19 | 45,698,914 | *QPCTL* | 45692400 | 45703987 | -7 | *isoQC* | 0.75 |
|  |  |  | *GIPR* | 45668187 | 45683722 | -31 | *Dh44-R2* | 0.56 |

**Supplementary Table 3.** Comparison of the multi-assays screen of BMI GWAS genes presented here and published data by Banranski et al. based on triglyceride (TAG) levels. Only 16 *Drosophila* candidate genes studied were common between the two studies but with these genes the results are largely agreeable suggesting potential involvement in the control of body weight.

| Human Gene | Drosophila gene | Same siRNA line | Adult TAG (Baranski et al.; 46) | Neuronal RNAi score |
| --- | --- | --- | --- | --- |
| SBK1 | CG4945 | * | fatter | 0.72 |
| NRXN3 | Nrx-1 | * | fatter | 0.90 |
| SEC16B | Sec16 |  | fatter | 0.86 |
| NUDT3 | Aps | * | fatter | 0.76 |
| COL4A3BP | cert |  | fatter | 0.91 |
| SLC39A8 | Zip71B |  | WT | 0.63 |
| SKOR1 | fuss | * | WT | 0.51 |
| NEGR1 | Lac |  | WT | 0.71 |
| TMEM18 | tmem18 |  | WT | 0.56 |
| LRP1B | LRP1 |  | WT | 0.76 |
| ATXN2L | Atx2 | * | WT | 0.97 |
| QPCTL | isoQC |  | WT | 0.75 |
| MTCH2 | Mtch |  | WT | 0.78 |
| KTCD15 | twz |  | WT | 0.70 |
| RPL27A | RpL27A |  | lethal | 0.89 |
| TUFM | EfTuM |  | lethal | 0.78 |

**Supplementary Table 4.** Genes that are reciprocally regulated in murine POMC and AgRP neurons in response to an overnight fast and their *Drosophila* orthologues.

| Mouse gene | POMC neurons |  |  | AgRP neurons |  |  | Drosophila orthologue |  |
| --- | --- | --- | --- | --- | --- | --- | --- | --- |
|  | Fed - Fast (TPM) | t test p value | Fold change | Fed - Fast (TPM) | t test p value | fold change | Gene name | % Identity |
| Orexigenic genes |  |  |  |  |  |  |  |  |
| **Arpc5l** | 62.6 | 0.021 | 1.5 | -54.5 | 0.013 | 1.5 | *Arpc5* | 44.0 |
| **Hsd17b12** | 35.9 | 0.002 | 2.0 | -46.3 | 0.000 | 2.1 | *spidey* | 44.0 |
| **Tubb4b** | 97.7 | 0.022 | 1.6 | -112.7 | 0.001 | 1.6 | *betaTub56D* | 94.0 |
| **Chml** | 1.8 | 0.041 | 2.6 | -3.6 | 0.025 | 1.6 | *Rep* | 33.0 |
| **Ddah1** | 94.4 | 0.007 | 1.8 | -79.1 | 0.001 | 2.6 | *CG1764* | 38.0 |
| **Fam162a** | 42.9 | 0.016 | 1.9 | -65.0 | 0.035 | 2.0 | *CG9231* | 32.0 |
| **Gem** | 19.7 | 0.036 | 2.3 | -74.9 | 0.000 | 4.9 | *Rgk1* | 30.0 |
| **Hcrt2** | 15.3 | 0.005 | 3.1 | -11.7 | 0.005 | 6.6 | *SIFaR* | 30.0 |
| **Prkcsh** | 30.0 | 0.048 | 1.5 | -49.6 | 0.045 | 1.7 | *GCS2beta* | 38.0 |
| **Ptplad1** | 132.2 | 0.034 | 1.5 | -170.7 | 0.000 | 1.6 | *CG9267* | 34.0 |
| **Spred2** | 40.5 | 0.033 | 2.4 | -69.3 | 0.002 | 1.9 | *Spred* | 35.0 |
| **Spred3** | 46.1 | 0.003 | 6.1 | -70.0 | 0.000 | 2.7 |  | 30.0 |
| **Ssr4** | 122.4 | 0.016 | 2.0 | -101.1 | 0.013 | 1.6 | *Tapdelta* | 34.0 |
| Anorexigenic genes |  |  |  |  |  |  |  |  |
| **Acvr2a** | 58.5 | 0.010 | 1.7 | -31.9 | 0.009 | 1.5 | *put* | 47.0 |
| **Aldh6a1** | 35.0 | 0.004 | 2.2 | -15.3 | 0.037 | 1.5 | *CG17896* | 67.0 |
| **Atp2b2** | 42.7 | 0.000 | 1.5 | -50.5 | 0.001 | 1.7 | *PMCA* | 60.0 |
| **Cacnb4** | 12.2 | 0.000 | 1.5 | -36.3 | 0.000 | 2.3 | *Ca-beta* | 53.0 |
| **Cdk14** | 30.7 | 0.016 | 1.6 | -12.8 | 0.002 | 4.6 | *Eip63E* | 52.0 |
| **Dtna** | 105.9 | 0.000 | 1.9 | -133.2 | 0.000 | 2.2 | *Dyb* | 43.0 |
| **Elavl2** | 14.0 | 0.006 | 1.5 | -32.8 | 0.002 | 2.0 | *Rbp9* | 58.0 |
|  |  |  |  |  |  |  | *fne* | 59.0 |
| **Epb4.1l4b** | 13.1 | 0.005 | 1.9 | -22.1 | 0.000 | 4.2 | *yrt* | 47.0 |
| **Gda** | 38.2 | 0.004 | 1.7 | -21.2 | 0.035 | 2.1 | *DhpD* | 45.0 |
| **Gpn3** | 51.6 | 0.001 | 2.9 | -24.1 | 0.006 | 1.8 | *CG2656* | 51.0 |
| **Nalcn** | 58.6 | 0.002 | 1.7 | -49.5 | 0.001 | 1.6 | *na* | 57.0 |
| **Nedd4l** | 33.6 | 0.010 | 1.7 | -35.5 | 0.000 | 1.7 | *Nedd4* | 49.0 |
| **Nmt1** | 20.8 | 0.006 | 1.5 | -21.9 | 0.044 | 1.5 | *Nmt* | 56.0 |
| **Pofut1** | 7.6 | 0.001 | 2.4 | -6.2 | 0.029 | 3.4 | *O-fut1* | 42.0 |
| **Prkaa1** | 23.4 | 0.028 | 1.5 | -21.3 | 0.019 | 1.5 | *AMPKalpha* | 60.0 |
| **Pura** | 83.5 | 0.021 | 1.5 | -111.8 | 0.014 | 1.9 | *Pur-alpha* | 44.0 |
| **Pygl** | 11.5 | 0.028 | 1.8 | -15.1 | 0.000 | 2.8 | *GlyP* | 71.0 |
| **Rasgef1a** | 29.8 | 0.005 | 1.5 | -15.6 | 0.031 | 1.5 | *CG7369* | 41.0 |
| **Rasgef1c** | 5.4 | 0.047 | 1.9 | -9.6 | 0.008 | 9.5 |  | 46.0 |
| **Rasgef1a** | 29.8 | 0.005 | 1.5 | -15.6 | 0.031 | 1.5 | *CG4853* | 36.0 |
| **Rasgef1c** | 5.4 | 0.047 | 1.9 | -9.6 | 0.008 | 9.5 |  | 37.0 |
| **Rps6ka6** | 12.7 | 0.028 | 2.1 | -40.1 | 0.014 | 2.4 | *S6kII* | 49.0 |
| **Rufy2** | 34.1 | 0.017 | 1.5 | -39.3 | 0.028 | 1.7 | *CG31064* | 42.0 |
| **Slc6a1** | 25.6 | 0.026 | 1.6 | -58.5 | 0.005 | 3.5 | *Gat* | 55.0 |
| **Syt7** | 4.3 | 0.036 | 2.9 | -13.9 | 0.008 | 1.9 | *Syt7* | 46.0 |
| **Tub** | 97.9 | 0.008 | 1.6 | -125.9 | 0.000 | 2.2 | *ktub* | 42.0 |
| **Ccdc132** | 20.5 | 0.047 | 1.6 | -16.8 | 0.010 | 1.5 | *Vps50* | 31.0 |
| **Fat3** | 15.7 | 0.016 | 2.0 | -28.3 | 0.002 | 6.7 | *kug* | 36.0 |
| **Hs6st2** | 93.9 | 0.003 | 1.7 | -35.4 | 0.022 | 1.5 | *Hs6st* | 38.0 |
| **Htr1a** | 12.1 | 0.035 | 5.4 | -30.9 | 0.000 | 4.9 | *5-HT1A* | 36.0 |
|  |  |  |  |  |  |  | *5-HT1B* | 35.0 |
| **Kcnj3** | 37.5 | 0.043 | 1.5 | -18.6 | 0.034 | 2.1 | *Irk2* | 35.0 |
| **Kcnq3** | 90.7 | 0.007 | 3.1 | -54.3 | 0.026 | 2.5 | *KCNQ* | 31.0 |
| **Lipa** | 26.0 | 0.006 | 1.6 | -19.9 | 0.000 | 2.4 | *CG18301* | 34.0 |
| **Lrch2** | 36.5 | 0.001 | 1.5 | -48.6 | 0.026 | 1.8 | *Lrch* | 31.0 |
| **Nr1d2** | 67.2 | 0.002 | 2.8 | -66.8 | 0.000 | 2.3 | *Eip75B* | 30.0 |
| **Rbfox1** | 40.1 | 0.005 | 1.6 | -23.1 | 0.000 | 2.5 | *Rbfox1* | 36.0 |
| **Rps6ka5** | 22.1 | 0.001 | 2.7 | -31.3 | 0.000 | 2.2 | *JIL-1* | 38.0 |
| **Slc44a5** | 15.4 | 0.026 | 3.0 | -6.0 | 0.033 | 1.7 | *Ctl2* | 34.0 |
| **Smpdl3b** | 4.4 | 0.041 | 3.9 | -5.1 | 0.008 | 5.9 | *CG32052* | 32.0 |
| **Sytl4** | 11.0 | 0.033 | 2.4 | -6.9 | 0.000 | 3.0 | *btsz* | 35.0 |
| **Vps39** | 46.6 | 0.007 | 1.6 | -56.1 | 0.004 | 1.7 | *Vps39* | 36.0 |

**Supplementary Table 5.** Comparison between assays overall score with or without triglyceride (TAG) and in comparison to wet mass (mass). Weight mass is a good substitute for the triglyceride measurement that is ideal for high throughput screens.

| Fly Gene | Mouse Gene | cafe & overfed & TG & mass & glucose | cafe & overfed & TG & glucose | cafe & overfed & mass & glucose |
| --- | --- | --- | --- | --- |
| **<u>"Orexigenic" genes</u>** |  |  |  |  |
| betaTub56D | Tubb4b | 0.692547395 | 0.633369224 | 0.80966566 |
| SIFaR | Hcrt2 | 0.670888869 | 0.652900126 | 0.670775078 |
| CG9231 | Fam162a | 0.678086043 | 0.696881884 | 0.627272354 |
| spidey | Hsd17b12 | 0.585400911 | 0.614727898 | 0.59183505 |
| Tapdelta | Ssr4 | 0.526868847 | 0.480457873 | 0.563057866 |
| GCS2beta | Prkcsh | 0.502633663 | 0.402022672 | 0.508165781 |
| Arpc5 | Arpc5l | 0.452842786 | 0.551967883 | 0.411279229 |
| CG1764 | Ddah1 | 0.469415051 | 0.367140667 | 0.363419649 |
| Spred | Spred2 & Spred3 | 0.465022366 | 0.540949619 | 0.345923868 |
| Rgk1 | Gem | 0.418702236 | 0.505046401 | 0.28518198 |
| Rep | Chml | 0.39050888 | 0.333205746 | 0.283265249 |
| **<u>"anorexigenic" genes</u>** |  |  |  |  |
| Eip75B | Nr1d2 | 0.996770208 | 0.99999 | 0.99596526 |
| Ca-beta | Cacnb4 | 0.975701453 | 0.977523713 | 0.969705264 |
| Dyb | Dtna | 0.938481401 | 0.945749783 | 0.924555783 |
| Lrch | Lrch2 | 0.896280386 | 0.880706433 | 0.924265579 |
| ktub | Tub | 0.865312824 | 0.846508131 | 0.847184882 |
| O-fut1 | Pofut1 | 0.859818163 | 0.842028118 | 0.840406424 |
| DhpD | Gda | 0.858489367 | 0.831008605 | 0.826890997 |
| S6kII | Rps6ka6 | 0.840390767 | 0.805278039 | 0.801553356 |
| Nedd4 | Nedd4l | 0.832924609 | 0.791726789 | 0.800300762 |
| JIL-1 | Rps6ka5 | 0.81642222 | 0.792938496 | 0.783376457 |
| fne | Elavl2 | 0.790819371 | 0.760255878 | 0.754351965 |
| Rbp9 | Elavl2 | 0.629212499 | 0.616160212 | 0.719379868 |
| CG31064 | Rufy2 | 0.720549997 | 0.66404147 | 0.717047961 |
| 5-HT1A | Htr1a | 0.768161105 | 0.787773521 | 0.712602463 |
| Irk2 | Kcnj3 | 0.703660335 | 0.642517335 | 0.684209878 |
| Syt7 | Syt7 | 0.592934481 | 0.511611567 | 0.680675656 |
| btsz | Sytl4 | 0.711426298 | 0.751904526 | 0.671894004 |
| CG7369 | Rasgef1a & Rasgef1c | 0.729906399 | 0.784360954 | 0.66352587 |
| CG17896 | Aldh6a1 | 0.712436037 | 0.775408126 | 0.640945046 |
| Ctl2 | Slc44a5 | 0.707615386 | 0.790276428 | 0.639480154 |
| GlyP | Pygl | 0.567213887 | 0.545724073 | 0.616067813 |
| 5-HT1B | Htr1a | 0.673426426 | 0.599774271 | 0.614702251 |
| yrt | Epb4.1l4b | 0.628113513 | 0.617693043 | 0.589905289 |
| Rbfox1 | Rbfox1 | 0.670455881 | 0.7383 | 0.588069851 |
| Eip63E | Cdk14 | 0.666701928 | 0.716463214 | 0.586494208 |
| CG32052 | Smpdl3b | 0.490245508 | 0.455160086 | 0.553117407 |
| Pur-alpha | Pura | 0.474793126 | 0.414616727 | 0.519214266 |
| Vps50 | Ccdc132 | 0.54735124 | 0.507738785 | 0.51655354 |
| CG18301 | Lipa | 0.493379832 | 0.451887594 | 0.510649465 |
| put | Acvr2a | 0.557258722 | 0.482107785 | 0.47449481 |
| Vps39 | Vps39 | 0.50003187 | 0.517006217 | 0.463701461 |
| CG2656 | Gpn3 | 0.492629776 | 0.580540686 | 0.460641316 |
| KCNQ | Kcnq3 | 0.52725463 | 0.553993136 | 0.422457285 |
| PMCA | Atp2b2 | 0.414415563 | 0.493272874 | 0.413822912 |
| CG4853 | Rasgef1a & Rasgef1c | 0.359659983 | 0.317208984 | 0.400421899 |
| na | Nalcn | 0.356676071 | 0.302911592 | 0.396054475 |
| kug | Fat3 | 0.343047534 | 0.297800378 | 0.355293841 |
| Nmt | Nmt1 | 0.365093596 | 0.361249656 | 0.312621303 |
| AMPKalpha | Prkaa1 | 0.377350758 | 0.377751294 | 0.245189833 |

**Supplementary Table 6.**
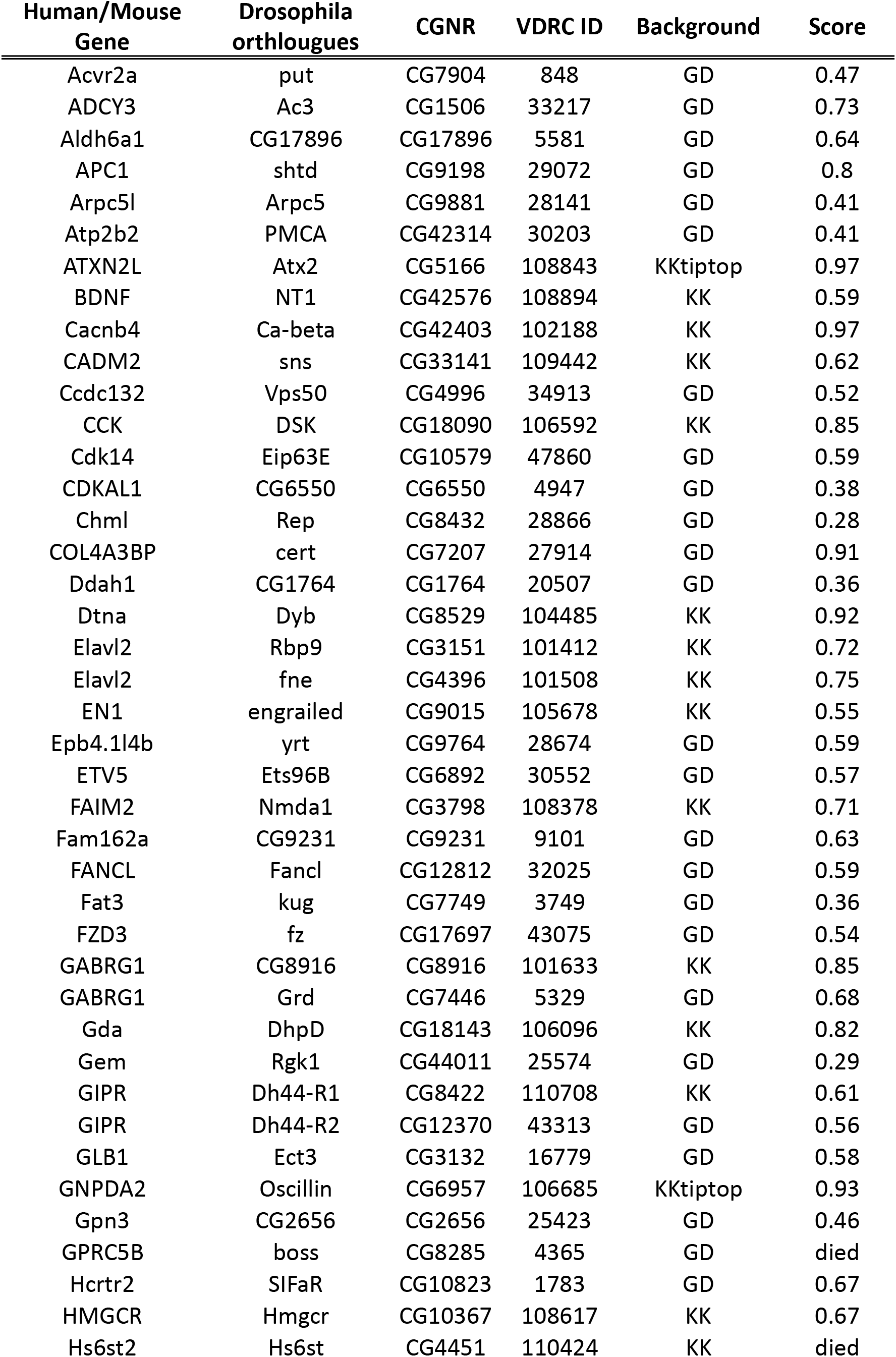
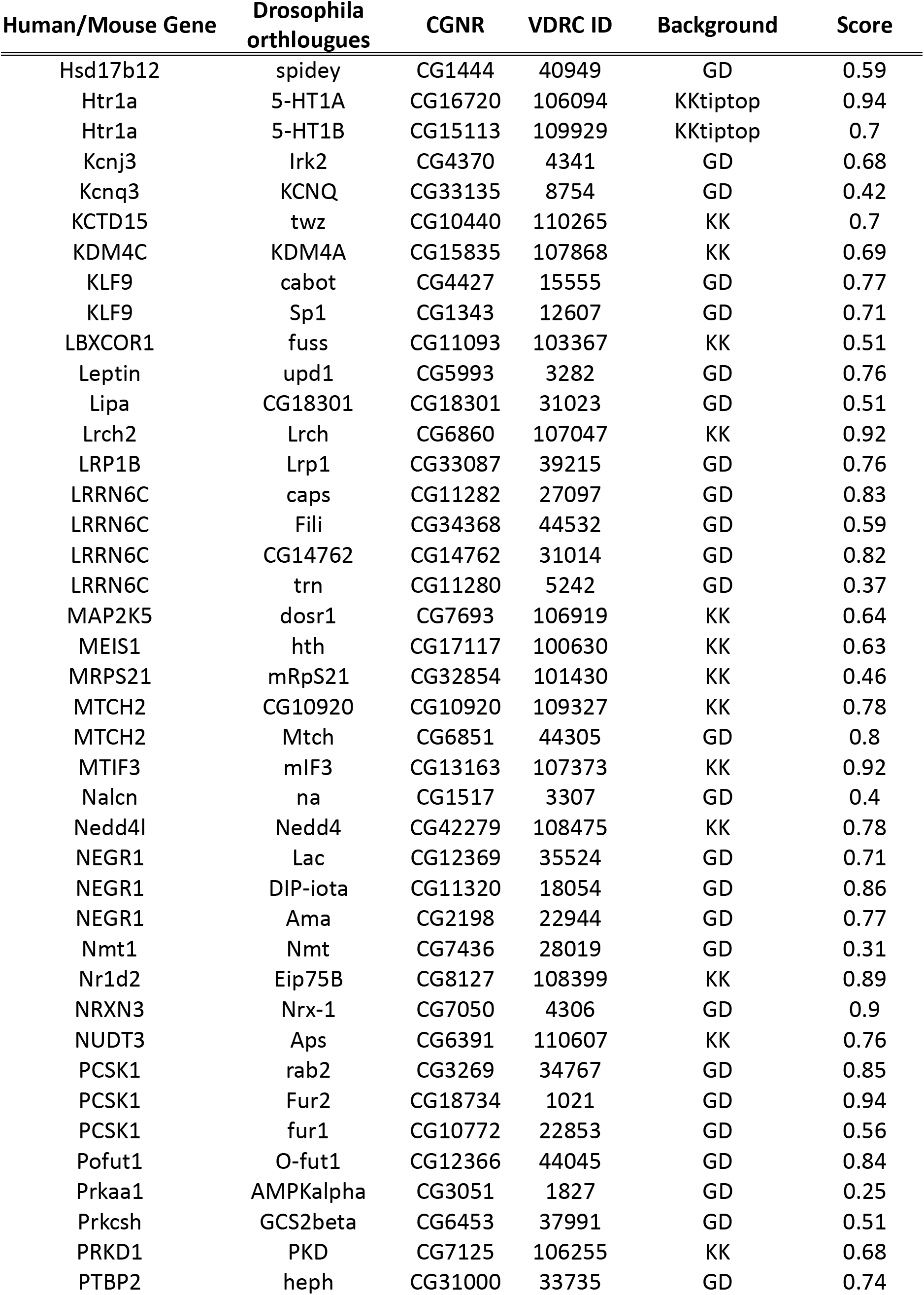
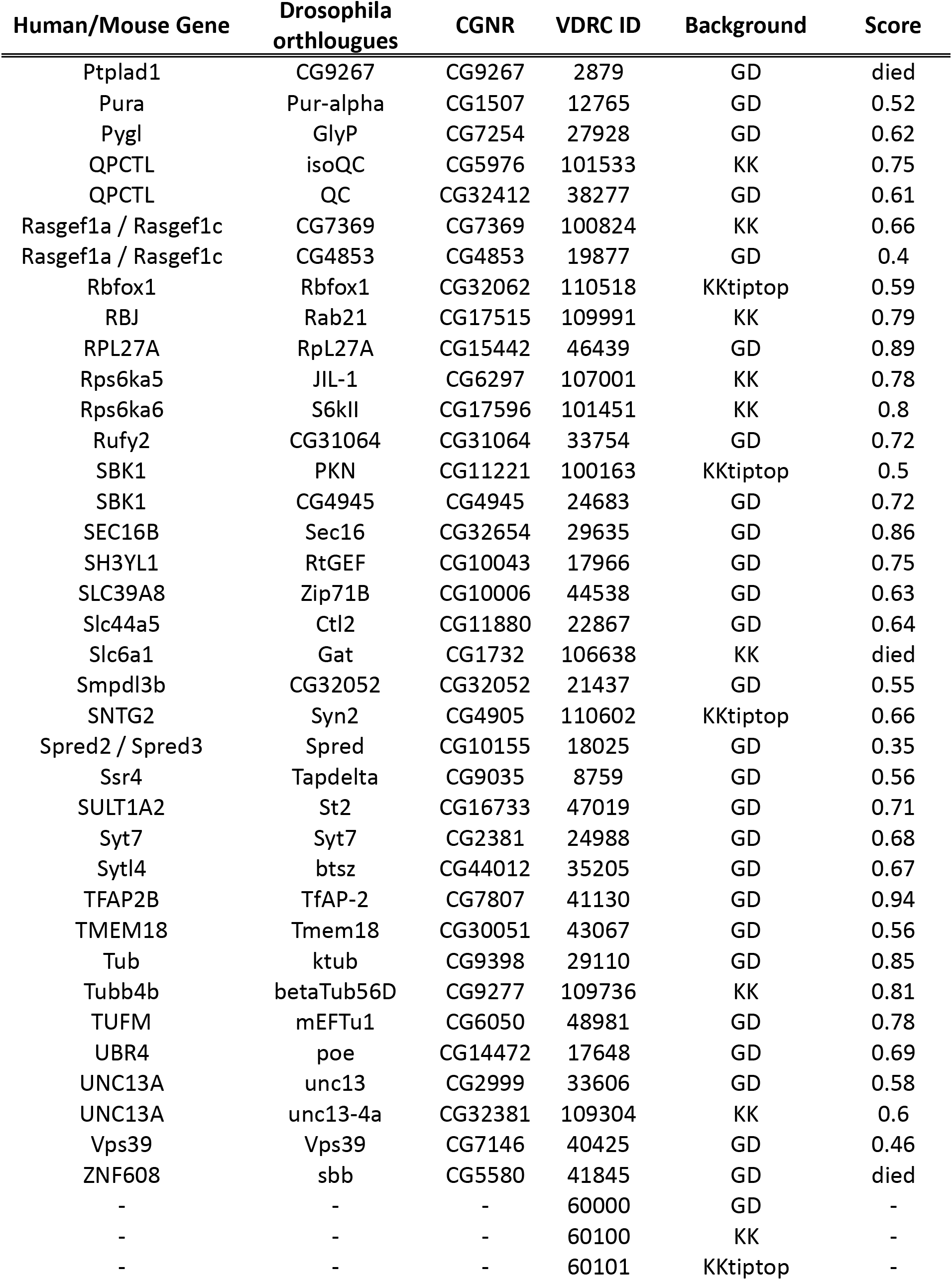
List of fly stocks used in this study that were purchased from the Vienna *Drosophila* Resource Center (VDRC).

## Acknowledgments

We thank the University of Cambridge fly facility for their continuous support in maintaining our fly stocks and the custom made fly foods. We thank the Cambridge Core Biochemical Assay Laboratory for the glucose and triglyceride assays. We thank the Vienna *Drosophila* Stock Centres for stocks.

